# Spatial RNA sequencing identifies robust markers of vulnerable and resistant human midbrain dopamine neurons and their expression in Parkinson’s Disease

**DOI:** 10.1101/334417

**Authors:** Julio Aguila, Shangli Cheng, Nigel Kee, Ming Cao, Menghan Wang, Qiaolin Deng, Eva Hedlund

## Abstract

Defining transcriptional profiles of substantia nigra pars compacta (SNc) and ventral tegmental area (VTA) dopamine neurons is critical to understanding their differential vulnerability in Parkinson’s Disease (PD). Here, we determine transcriptomes of human SNc and VTA dopamine neurons using LCM-seq on a large sample cohort. We apply a bootstrapping strategy as sample input to DESeq2 and identify 33 stably differentially expressed genes (DEGs) between these two subpopulations. We also compute a minimal sample size for identification of stable DEGs, which highlights why previous reported profiles from small sample sizes display extensive variability. Network analysis reveal gene interactions unique to each subpopulation and highlight differences in regulation of mitochondrial stability, apoptosis, neuronal survival, cytoskeleton regulation, extracellular matrix modulation and well as synapse integrity, which could explain the relative resilience of VTA dopamine neurons. Analysis of PD tissues showed that while identified stable DEGs can distinguish the subpopulations also in disease, the SNc markers SLIT1 and ATP2A3 were downregulated and thus appears to be biomarkers of disease. In summary, our study identifies human SNc and VTA marker profiles, which will be instrumental for studies aiming to modulate dopamine neuron resilience and to validate cell identity of stem cell-derived dopamine neurons.

## INTRODUCTION

Midbrain dopamine neurons are divided into two major populations, the substantia nigra pars compacta (SNc) and the ventral tegmental area (VTA) [22]. SNc dopamine neurons project to the dorsolateral striatum [12] and are severely affected in Parkinson’s Disease (PD) [13,14], while VTA dopamine neurons project to cortical and mesolimbic areas and are more resilient to degeneration [22]. These neuron populations have been extensively investigated in numerous rodent models, [19,11,18,6,45], towards the goal of identifying molecular mechanisms that can prevent degeneration or to model disease. Targeted analysis of midbrain dopamine neuron populations has revealed several markers that appear to differentially label SNc e.g. *Aldh1a7, Sox6, Cbln1, Vav3, Atp2a3* and VTA e.g. *Calb1, Otx2, Crym, Cadm1* and *Marcks* [13,19,11,18,15,6,41,39]. Transcriptional analysis of human tissue has largely been limited to SNc [8,50] except for our recent small sample cohort to compare SNc and VTA [39]. These aforementioned investigations display extensive cross-study variability, resulting in very few reproducible markers either within mouse, rat and human or across different species. Small sample sizes could be a confounding factor of these studies, along with differences in rodent strain backgrounds, methodological differences, or inter-individual variability among human patients.

To reveal cell intrinsic properties underlying the differential vulnerability of SNc and VTA dopamine neurons in PD, a thorough large-scale transcriptional profiling in adult human tissues is required. Such an analysis could also investigate the necessary minimum cohort size, above which lineage specific markers remain stably differentially expressed irrespective of patient selection, an essential requirement for valid study design in variable human populations. Finally, identified differences could also serve as a foundation for the selective *in vitro* derivation of SNc dopamine neurons, which represent the ideal cell type for transplantation in PD [50,21,55,22,29,17].

Here we used the spatial transcriptomics method LCM-seq, which combines laser capture microdissection with Smart-seq2 RNA sequencing [38,39], to precisely analyse individually isolated SNc and VTA dopamine neurons from 18 human *post-mortem* brains. Using bootstrapping without replacement coupled with DESeq2, we identify 33 markers that were stably differentially expressed between SNc and VTA dopamine neurons. We show that the minimal sample size required to reliably identify these subtype-specific markers in this cohort is eight subjects, which may explain why smaller cohorts have given inconsistent results. Several of the markers identified here have been implicated in PD or other degenerative diseases and thus provide compelling future targets to modulate neuronal vulnerability or to model disease. We also analysed the regulation of these stable genes in PD patient tissues and found that these markers still define the two subpopulations in end-stage disease and that only two SNc markers, SLIT1 and ATP2A3, were severely down-regulated in PD.

## MATERIALS AND METHODS

### Ethics statement

We have ethical approval to work with human *post-mortem* samples (Supplemental Tables S2 and S3) from the regional ethical review board of Stockholm, Sweden (EPN Dnr 2012/111-31/1; 2012/2091-32). Fresh frozen tissue was obtained through the Netherlands Brain Bank (NBB). The work with human tissues was carried out according to the Code of Ethics of the World Medical Association (Declaration of Helsinki).

### Tissue sectioning and laser capture

Sample preparation prior LCM-seq was carried out as follows. Frozen midbrain tissues (controls and PD) obtained from the brain banks were attached to chucks using pre-cooled OCT embedding medium (Histolab). 10 μm-thick coronal sections were acquired in a cryostat at −20 °C and placed onto precooled-PEN membrane glass slides (Zeiss). For RNAscope experiments (control tissue), sections were cut at 12 μm-thickness and attached to Superfrost^®^ Plus slides (Thermo Scientific). The slides with sections were kept at −20 °C during the sectioning and subsequently stored at −80 °C until further processed. The laser capture procedure followed by sequencing library preparation (LCM-seq) was carried out as described [38,39].

### Mapping and gene expression quantification

Samples were sequenced using an Illumina HiSeq2000, HiSeq2500 or NovaSeq platforms (reads of 43 or 50 bp in length). The uniquely mapped reads were obtained by mapping to the human reference genome hg38/GRCh38 using STAR with default settings. The reads per kilobase of transcript per million mapped reads (RPKM) were estimated using “rpkmforgenes” [45] to 10.88 million reads and 4.7 to 12.3 thousand genes expressed with RPKM>1, all samples were included. For control subjects the correlation coefficient between any two nearest samples was above 0.7. For PD samples we verified that all samples had > 1 million reads > 4600 genes expressed with RPKM >1. For PD samples the correlation coefficient between any two nearest samples was above 0.9. It should be noted that it has been elegantly demonstrated that shallow RNA sequencing of ca 50,000 reads/cell is sufficient for unbiased cell type classification and marker gene identification of neural subclasses [44] and thus our sequencing depth of >1 million reads/sample should be more than sufficient to subclassify SNc and VTA dopamine neurons. For all control or PD cases having more than one replicate per group, corresponding samples were averaged before analysis so that each case had only one SNc and one VTA. We confirmed the expression of known midbrain dopamine neuron markers and the purity of each sample (Fig. 1A and Supplemental Fig. S4).

**Figure 1.**
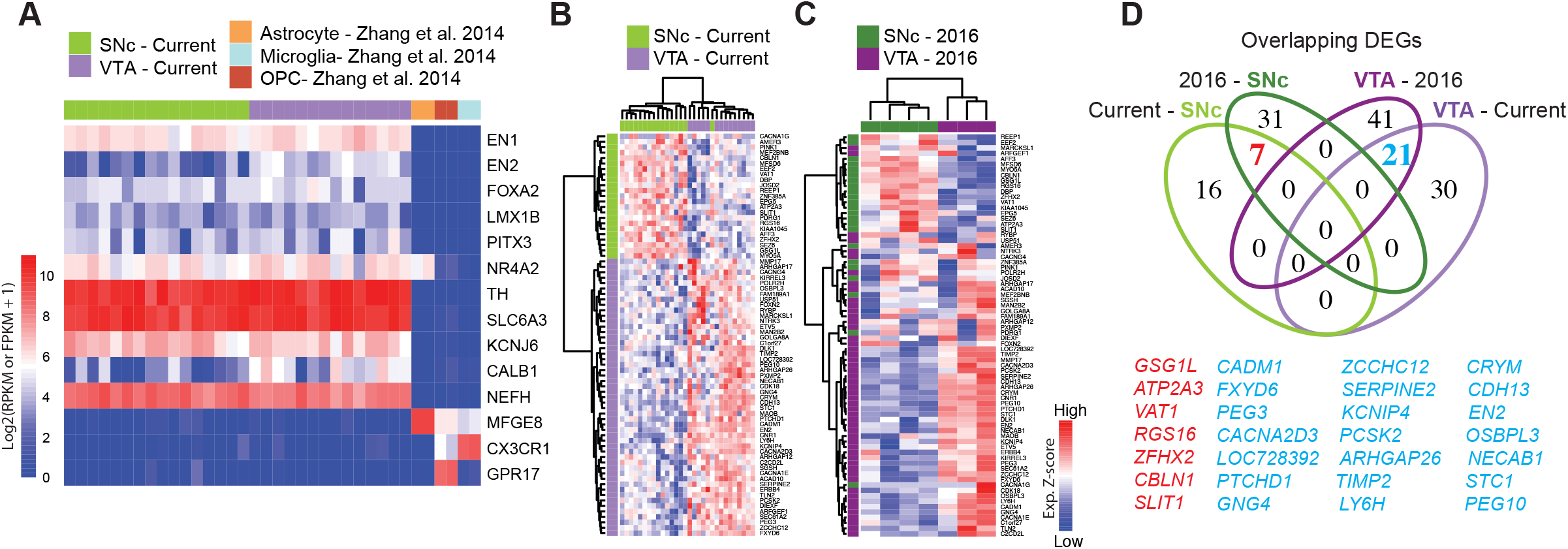
Gene signature of human adult midbrain dopamine neurons using the spatial sequencing method LCM-seq. (A) The high sample quality for the 18 male subjects profiled in this study was confirmed by strong expression of the midbrain dopamine neuron markers *EN1/2, FOXA2, LMX1B, PITX3, NR4A2, TH* and *SLC6A3* (DAT), the pan-neuronal marker neurofilament (*NEFH*), and the lack of astrocyte, microglia or oligodendrocyte precursor marker [61] contamination. (B) Hierarchical clustering analysis of samples from the current study using the 74 DEGs identified by DESeq2. (C) The 74 DEGs also separated SNc and VTA samples from [39] (3 female subjects). (D) Venn-diagram showing the relatively low degree of overlap between DEGs in the cohorts of different sizes. See also Supplemental Fig. S2.

### Differential expression analyses

Differentially expressed genes were identified using the R package “DESeq2” (version: 1.16.1) [33] where the cutoff for significance was an adjusted P value of 0.05. Identified DEGs (from different analysis and summarized below) are shown in Supplemental Tables S4 and S6 to S9.

Supplemental Table S6: Human differentially expressed genes (current study): 74 DEGs calculated from 16 SNc and 14 VTA samples from 18 control male individuals.

Supplemental Table S7: Human differentially expressed genes [39]: 100 DEGs calculated from 4 SNc and 3 VTA samples from 3 female subjects.

Supplemental Table S8: Human stable genes: 33 DEGs calculated from 12 control individuals (from the current study) using the bootstrapping approach.

### Bootstrapping approach coupled with DESeq2

To counteract the variability among human subjects and identify the most reliable DEGs between SNc and VTA neurons across datasets we developed a bootstrapping approach coupled with DESeq2 (Fig. 2A; Supplemental Fig. S3A). The stable genes output of this analysis is correlated with the sample size and give an unbiased estimation of the number of individuals required to consistently distinguish these closely related subpopulations. Importantly this computational tool can be used for the comparison of any other two groups, ().

**Figure 2.**
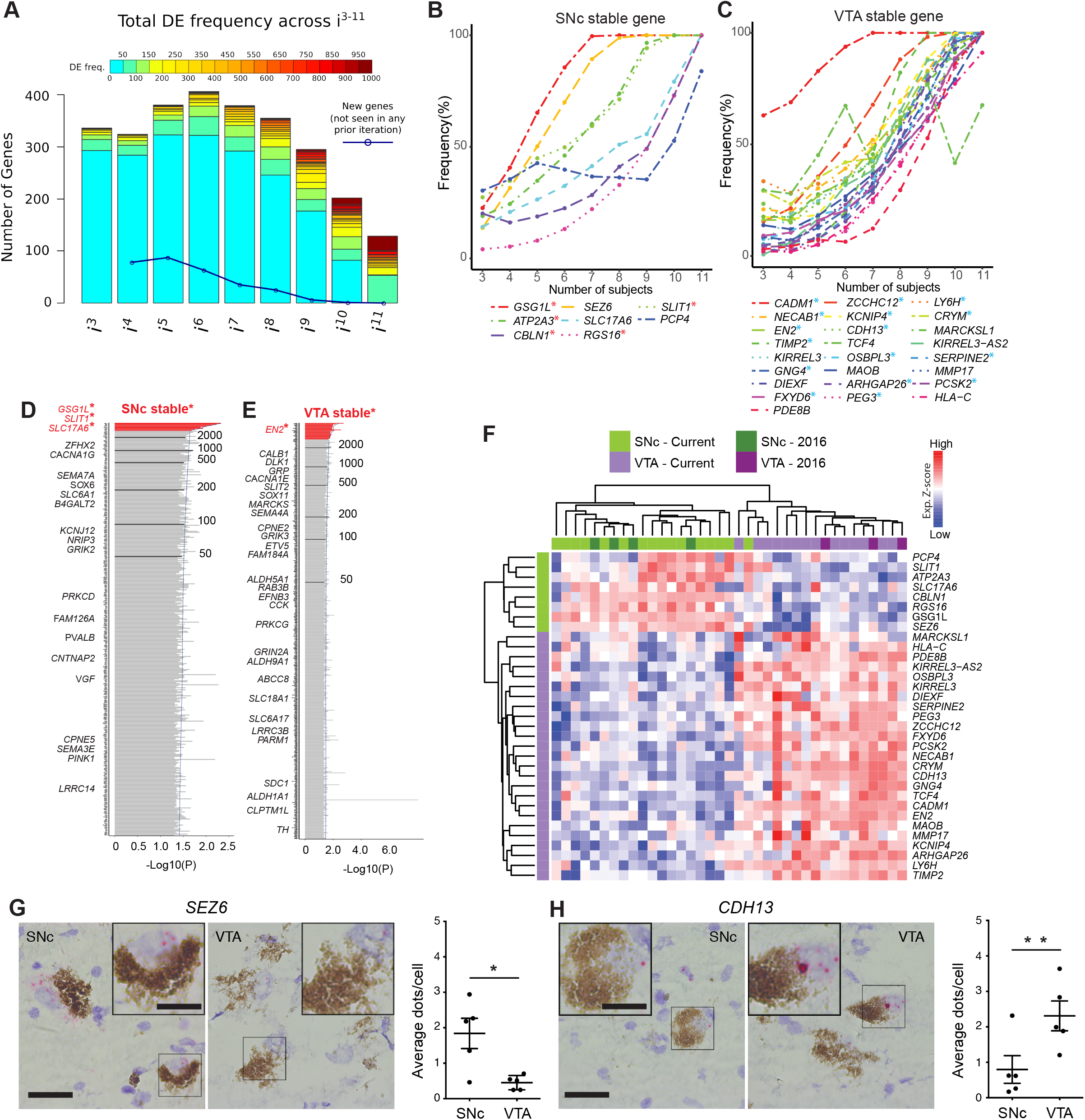
Bootstrapping analysis coupled with DESeq2 identifies SNc or VTA stable genes from a cohort of 12 human subjects. (A) Histogram of DEG frequency through iterative bootstrapping. X axis denotes increasing size of patient pool (i^3^-i^11^ individuals) at each iteration. Y-axis bar height denotes the number of DE genes, while bar color denotes the frequency those genes being DE in that iteration. Blue line denotes the decreasing number of novel genes detected across successive iterations. (B,C) DE frequency increases at each iterative sample size increase, for SNc (C) or VTA (D) stable genes. (D,E) Frequency histograms (ranked by p-value), output of the bootstrapping approach, extend the analysis of identified SNc and VTA stable genes. Genes with the highest frequency (at the top, in red) represent SNc (C) and VTA (D) stable genes. (F) Hierarchical clustering analysis using the 33 stable genes, faithfully segregates both SNc and VTA samples. (G,H) RNAscope staining and quantification for the stable genes SEZ6 (F, p=0.025) and CDH13 (G, p=0.005) enriched in the SNc and VTA, respectively (n=5 subjects, data represented as mean ± SEM, Paired t test). Scale bars 30 μm (15 μm for insets)

In detail:

1. Define ^..^N^..^ and ^..^M^..^ as the number of samples in Group 1 and Group 2, respectively. Choose ^..^I^..^ as a reference representing a given number of samples from ^..^N^..^ and ^..^M^..^.
2. Define ^..^i^..^ as the number of randomly selected samples from Group1 and Group2, where i ∈{3, 4, 5, …, ^..^I-1^..^}. In the human dataset, as we have 12 paired samples, the i ∈{3, 4, 5, 6, 7, 8, 9, 10, 11}.
3. Pool ^..^i^..^ samples (temporary considered a ^..^new data set^..^) and calculate DEGs with DESeq2.
4. Repeat steps 2) and 3) for ^..^j^..^ times (set to 1000 times in this study).
5. For every round of random selection and DESeq2, save the full list of DEGs, compute and rank their frequency.
6. Set a threshold (30% ratio in this study) to consider DEGs with higher frequency as stable genes.

Reliable genes appear when frequencies are above: Total times of bootstrapping x ratio (300 in this study). A stringent, but fair ratio can be defined by comparing the percentage of identified stable genes overlapping with the top (most significant) 10%, 20%, 30%, …, DEGs identified by DESeq2 alone.

### Bootstrapping approach applied to human samples

To reliable identify DEGs between human SNc and VTA samples, while minimizing subject variability, we selected 12 control individuals (66% of the dataset, 12 out of 18 individuals) where both neuronal populations were available and sequenced. Hence, the number of randomly selected samples (^..^n^..^ and ^..^m^..^ from ^..^i^..^ individuals) was from three to 11 and the algorithm repeated 1000 times (Fig. S3AB).

### Bootstrapping applied to mouse single cells

For this adult mouse dataset [30] we defined the groups SNc (N=73 cells) and VTA (M=170 cells comprising VTA1, VTA2, VTA3 and VTA4). To compensate the unbalance in cell number and adjust dataset representation compared to the human analysis (66%), we first randomly collected a subset of 73 VTA cells, pairing both SNc and VTA. Similarly, the number of randomly selected samples was 20, 25, 30, …, 70 and the algorithm repeated again 1000 times.

### STRING network based on DE genes between SNc and VTA

Based on the DE genes between SNc and VTA (*Supplementary Table S6*), two STRING networks were created separately. The MLC clustering grouped the network into sub-networks with default parameters and marked as the dash line.

### Data visualization

Data visualization was achieved using Principal Component Analysis (PCA) and Hierarchical Clustering (H-cluster). PCA was calculated with the function “prcomp” in R with default parameters. Then samples are projected onto the first two dimensions, PC1 and PC2. For H-cluster we used the R function “pheatmap” (version 1.0.12) with the clustering method of “ward.D2”.

### RNAscope staining of human tissues

RNAscope [58] was used to verify the expression (in control tissues) of one SNc marker (*SEZ6*) and one VTA-preferential gene (*CDH13*) based on the sequencing data. In brief, midbrain sections of human fresh frozen tissue (Supplemental Table S3) were quickly thawed and fixed with fresh PFA (4% in PBS) for 1 hour at 4’C. The RNAscope 2.5 HD Assay - RED Kit (Cat. 322360) was used using manufacturer recommendations. To evaluate the procedure in the midbrain tissue (Supplemental Fig. S3G), we first tested a negative control probe against a bacterial gene (Cat. 310043, *dapB*-C1) and a positive control probe against tyrosine hydroxylase (Cat. 441651, *TH*-C1) (Supplemental Fig. S3G). Once we set up the assay, midbrain sections were stained with *SEZ6* (Cat. 411351-C1) or *CDH13* probes (Cat. 470011-C1). Slides were counterstained with fresh 50% Gill Solution (Cat. GSH132-1L, Sigma-Aldrich) for 2 minutes, washed in water and dried for 15min at 60°C before mounting with Pertex (Cat. 00811, Histolab). For every sample (n=5), we imaged 5-6 random fields within the SNc and VTA regions. On average 194.25±43.02 cells were imaged per region and staining. Pictures were made at 40X magnification using the bright-field of a Leica microscope (DM6000/CTR6500 and DFC310 FX camera). After randomization and coding of all the images, the number of dots within melanised cells (dopamine neurons) were counted using ImageJ (version 1.48) and later the average number of dots per cells determined for each region. Investigators performing the quantification were blinded to the sample, target region (SNc and VTA) and probe staining.

### Statistical analysis

For this study, statistical analyses were performed using ^..^R^..^. For the RNAscope analysis a paired t test (Prism 6, Version 6.0f) was used to compare the mean average dots per cell (for *SEZ6* or *CDH13* staining) between the SNc and VTA. Where applicable, individual statistical tests are detailed in the figure legends where significance is marked by P < 0.05. The number of subjects/cells used for each experiment is listed in the figure or figure legends. Results are expressed as mean ± SD or SEM as specified in the figure legend.

### Data access

All raw and processed sequencing data generated in this study have been submitted to the NCBI Gene Expression Omnibus (GEO; https://www.ncbi.nlm.nih.gov/geo/) under accession number GSE114918. Human samples re-analysed from the Nichterwitz study [39]. ArrayExpress (E-MEXP-1416) [8] or raw data received from Dr. Kai C. Sonntag [51]. A processed table with RPKMs values for the full dataset generated in this study can also be found at Mendeley Data under DOI 10.17632/b7nh33pdmg.1

## RESULTS

### Published SNc and VTA transcriptional profiles display considerable discrepancies

To understand the molecular underpinnings of the differential vulnerability among dopamine neurons, we compared previously published transcriptome studies of mouse and rat VTA and SNc dopamine neurons, using the list of markers reported as significantly up- or down-regulated [19,11,18,6]. This analysis revealed that a surprisingly low fraction of DEGs were common across data sets (Supplemental Fig. S1A, B; Supplemental Tables S1 and S4). Comparing across species with our previously published small data set on human SNc and VTA [39], only two genes, *SOX6* and *CALB1*, overlapped within SNc and VTA gene lists, respectively (Supplemental Fig. S1C). These discrepancies highlight the urgent need to identify reproducible marker profiles for VTA and SNc dopamine neurons.

### LCM-seq of a large human cohort identifies markers specific to SNc or VTA dopamine neurons and suggests that sample size impacts identification of DEGs

To identify robust and specific human dopamine neuron subpopulation markers, we isolated individual VTA and SNc neurons from *post-mortem* tissues from 18 adult individuals by LCM (Supplemental Fig. S2A-G; Supplemental Table S2) and conducted polyA-based RNA sequencing. This study represents the largest human data set profiling of SNc and VTA dopamine neurons to date. The quality of human fresh frozen tissues used may vary as a consequence of *post-mortem* interval (PMI), sample handling and preservation. Therefore, prior to conducting differential gene expression analysis we performed extensive quality control analysis to rule out undesired influences from sample processing (Supplemental Table S5). Randomly selected samples that exhibited different PMIs for VTA or SNc neurons displayed comparable cDNA quality (Supplemental Fig. S2H, I). Furthermore, while the total number of reads varied between individual samples, such variability was similarly distributed between SNc and VTA samples (Supplemental Fig. S2J). The number of detected genes did not correlate with either the age of the donor (Supplemental Fig. S2L), the PMI (Supplemental Fig. S2M) or the total number of reads (Supplemental Fig. S2K). Only the number of collected cells per sample modestly impacted gene detection (P=0.515) (Supplemental Fig. S2N). However, neither the number of collected cells nor the number of detected genes were significantly different between SNc and VTA neuron groups (Supplemental Fig. S2O, P) and thus should not affect DEG identification. Finally, we observed that all samples strongly expressed the dopamine neuron markers *EN1/2, FOXA2, LMX1B, PITX3, NR4A2, TH* and *SLC6A3* (DAT), and the general neuronal marker NEFH, while they lack glial markers *MFGE8, CX3CR1* or *GPR17*. This clearly demonstrates the selective enrichment of dopamine neurons using the LCM-seq methodology (Fig. 1A). *KCNJ6* (GIRK2) and *CALB1*, two genes often used to distinguish between SNc or VTA dopamine neurons, were also expressed (Fig. 1A), but could not, on their own, accurately differentiate our samples (Supplemental Fig. S2Q).

Differential expression analysis, considering these 18 individuals, identified 74 DEGs (Supplemental Table S6), which resolved SNc from VTA neurons (Fig. 1B). These genes also distinguished SNc and VTA samples in our previous small human cohort (N=3) [39] (Fig. 1C). However, relatively few DEGs overlapped with the current large cohort (N=18), even though the same experimental method was used. In fact, only seven SNc and 21 VTA DEGs overlapped across the current and the previous cohorts (Fig. 1D). Notably, the 100 DEGs identified in the small cohort (N=3) [39] (Supplemental Fig. S2R), failed to distinguish SNc and VTA in the current larger cohort of 18 subjects (Supplemental Fig. S2S; Supplemental Table S7). This suggests that small sample size prevents confident identification of DEGs.

### Bootstrapping coupled with DESeq2 identifies stable DEGs unique to human SNc or VTA

To evaluate how sample size may affect DEG detection, we used a bootstrapping algorithm in combination with DESeq2. To reduce the biological variability, we considered only those subjects for which both SNc and VTA samples were available (12/18 subjects, 24 samples in total). To begin with this approach, a subset of three subjects were randomly chosen from the pool of total 12 subjects. Differential expression analysis was then performed between the SNc and VTA samples of these subjects (DESeq2), and DEGs were selected with adj P-value < 0.05. This random sampling of three subjects, followed by DESeq2 analysis, was performed a total of 1,000 times, and the DE frequency over these 1,000 comparisons was recorded for this iteration (i = 3 subjects). Subsequently, this process was repeated using subsets of four subjects, then five, up to a maximum of 11 of the 12 subjects. For each subset size (i^3^ to i^11^) the DEG frequency was calculated by the 1,000x comparisons of that iteration (Supplemental Fig. S3A). By considering all DEGs in an iteration we were able to detect hundreds of genes, where on average only e.g. 13 DEGs were detected in i^3^ (Supplemental Fig. S3B). We found that the number of DEGs decreased with increased subset size, while the detection frequency increased. More importantly, the number of new DEGs detected also decreased with increasing sample size (Fig 2A). Interestingly, we found that in our cohort eight subjects were required to saturate the DEG detection, as few new genes were identified when considering additional subjects in subsequent iterations (Fig. 2A, blue line). We identified eight stable genes for the SNc (Fig. 2B) and 25 stable genes for the VTA (Fig. 2C). Five of the SNc stable genes (labeled in red**\***, see Fig 1D: *GSG1L, ATP2A3, CBLN1, RGS16* and *SLIT1*) and 16 of the VTA stable genes (in blue**\***, see Fig 1D: *CADM1, NECAB1, EN2, TIMP2, GNG4, FXYD6, ZCCHC12, KCNIP4, CDH13, OSBPL3, ARHGAP26, PEG3, LYH6, CRYM, SERPINE2* and *PCSK2*) were among the aforementioned seven and 21 overlapping DEGs, that we identified across the two studies.

We then summed DEGs across i^3^ to i^11^ (9,000 comparisons in total), separated genes into SNc or VTA enriched lists, and ranked the lists from most to least frequently DEs (Fig. 2D, E; Supplemental Fig. S3C, D; Supplemental Table S8). Highly ranked genes on these two lists included multiple known SNc and VTA markers in human (e.g. *GSG1L, SLIT1, ATP2A3, CADM1, CRYM, TCF4*) [8,50,30,39]. To identify the most reliable DEGs, we designated genes that were detected more than 3,000 times (out of 9,000) as “stable genes”. This stringent cutoff was chosen since the resulting SNc and VTA lists would then contain at least one stable gene that could be identified during the first iteration (where three individuals were used as the sample size). Frequency histograms (ranked by p-value) as the output of the overall bootstrapping approach showed genes with the 30% threshold (at the top, in red) represent SNc and VTA stable genes (Supplemental Fig. S3C). We also compared this stable gene list with the outcome of DESeq2 analysis alone, when applied to the same 12 subjects (Supplemental Fig. S3D, E). All eight SNc and 25 VTA markers perfectly overlapped with the DEGs from DESeq2 alone using an adjusted *P*-value <0.05 (Supplemental Fig. S3D) or a more stringent significance (adj. *P*-value <0.01, Supplemental Fig. S3E). The expression of SNc stable genes was confirmed in two independent human microarray datasets which only analyzed SNc neurons (Supplemental Fig. S3F [8,51]. Importantly, the stable genes faithfully classified SNc and VTA from 21 individuals (Fig. 2F), namely all 18 male individuals from our current dataset and the three female samples investigated previously [39]. Moreover, using RNA scope we confirmed the subpopulation-specific expression pattern of the identified SNc stable gene *SEZ6* (Fig. 2G) and the VTA stable gene *CDH13* (Fig. 2H) in human *post-mortem* tissues, further ratifying our LCM-seq data and the bootstrapping approach.

In conclusion, we have identified 33 markers that correctly classify samples as either SNc or VTA, and that remain robust to individual subject variability. Notably, these genes were stably differentially expressed only when at least eight subjects were included in the bootstrapping strategy (Fig. 2B, C). Thus, we have defined the minimal sample size required to distinguish SNc and VTA subpopulations in human subjects using LCM-seq and show that DEG identification below this number is unreliable. Depending on the variability among samples within a particular cohort this number could variate and should thus first be defined for each new cohort. The variability in DEGs identified between SNc and VTA dopamine neurons among previous studies could in part be explained by their use of too small cohorts.

To further validate our bootstrapping approach, we applied it to a published, postnatal, mouse single-cell dataset profiling midbrain dopamine neurons [30] (Supplemental Fig. S4A, raw data analyzed here). Single cells were initially assessed for expression of known dopamine neuron markers and the absence of contaminating glia or oligodendrocyte markers (Supplemental Fig. S4B) [61]. All available SNc dopamine neurons (73 in total) and 73 randomly selected VTA dopamine neurons were then subjected to aforementioned bootstrapping followed by DESeq2, through which we identified 36 SNc-enriched transcripts and 53 VTA-enriched transcripts (Supplemental Fig. S4C, D; Table S9). These stable gene sets for SNc and VTA included novel genes in addition to previously reported markers [6,11,18,19,30,39,45]. Importantly, these 89 stable genes, identified through our bootstrapping approach, effectively classified the single cells into the correct population, SNc or VTA, (Supplemental Fig. S4E) and the specific expression patterns in either SNc or VTA was corroborated in the adult mouse using Allen *in situ* images as exemplified in Supplemental Fig S4F. Specific expression patterns within either SNc or VTA was confirmed using Allen Brain Atlas, see examples of *Serpine2, Zcchc1* and *Cdh13* in coronal midbrain sections (Supplemental Fig. S4F). Finally, we wanted to see how the stable DEGs would overlap with DEGs identified through DEseq2 alone. We first plotted the number of DEGs identified through DESeq2 as a function of the adjusted P-value with an evident and expected decrease in the number of identified DEGs between SNc and VTA with stricter P-values (Supplemental Fig. S4G). We then plotted the stable SNc and VTA DEGs and the DEGs identified through DESeq2 alone at an adjusted P-value=0.05. The resulting Venn diagram shows that the majority of stable DEGs identified through our bootstrapping approach were also included when DESeq2 alone was used., with 50 out of 53 stable VTA DEGs and 25 out of 36 stable SNc DEGs being identified (Supplemental Fig. S4H).

In conclusion, with the bootstrapping strategy could be reliably applied to another larger data set and used to define stable SNc and VTA markers between two highly similar populations.

### STRING analysis identifies novel networks for human SNc and VTA dopamine neurons and highlights cellular functions that uniquely define each subpopulation

To explore potential interactions among the stable DEGs in human SNc and VTA dopamine neurons, we conducted STRING analysis. For this purpose, we used the 74 DEGs, including the stable DEGs, retrieved by comparing SNc and VTA, as the input (Supplemental Table S5). The two STRING networks shown are thus based on 23 genes with preferential expression in SNc (Fig. 3A) and 51 genes with predominant expression in VTA (Fig. 3B). The interactions between genes are shown through different color edges and were curated from databases, experiment or prediction. The nodes of the two networks are grouped using dashed lines by using MCL (Markov Clustering), in which the nodes with solid edges are from sub-networks. The interactions of the DEGs present in the VTA network highlight possible beneficial functions that are predominant in this resilient dopamine neuron subpopulation, including induction of survival genes, regulation of mitochondrial stability, catabolism of dopamine, regulation of resting membrane potential, extracellular matrix modulation and regulation of cytoskeleton and synapse integrity (Fig. 3C). The identified enriched gene networks give clues to networks that underlie the subpopulations unique functions and likely their differences in susceptibility.

**Figure 3.**
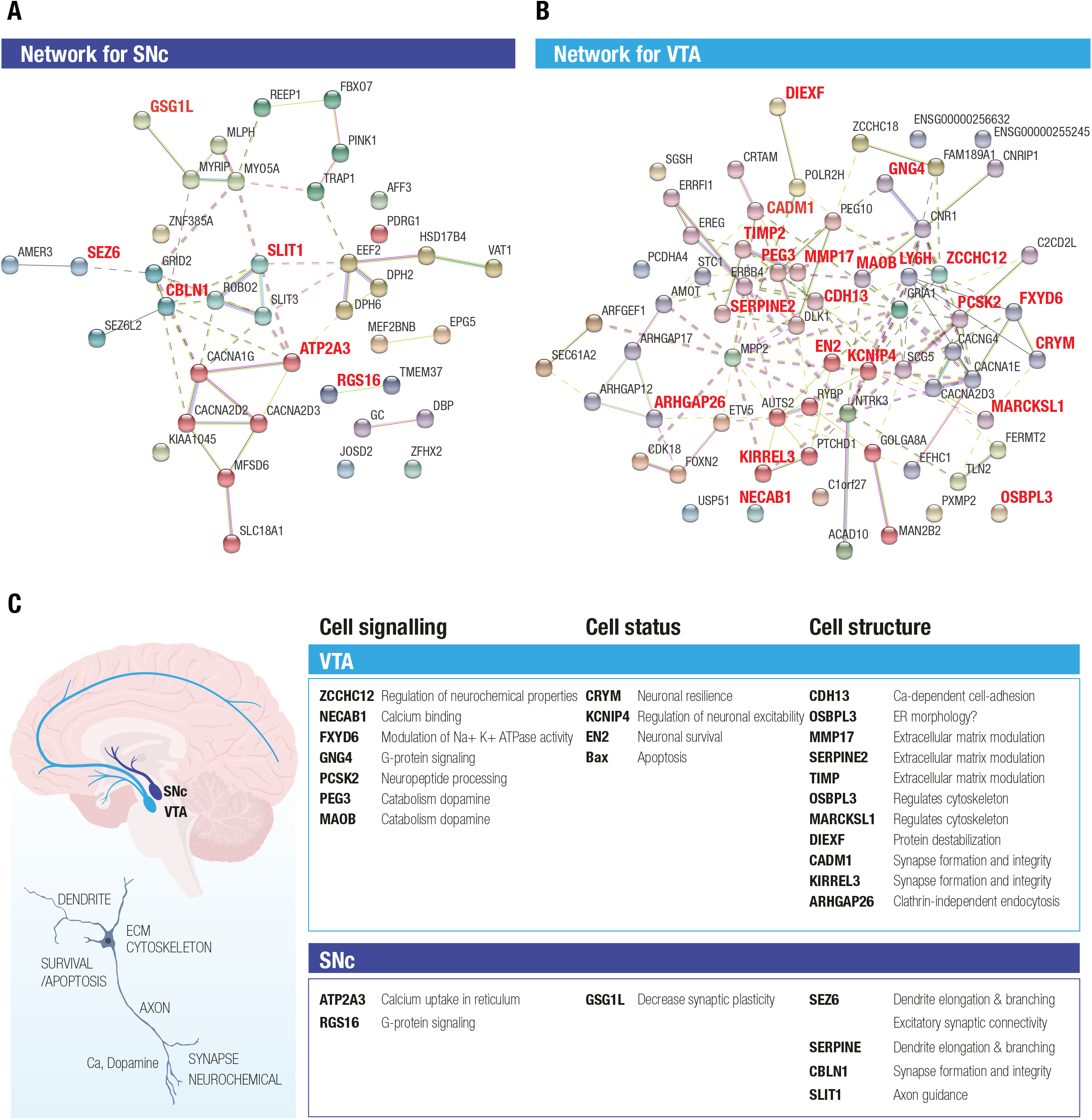
STRING analysis of DEGs between SNc and VTA identifies novel networks for the two midbrain dopamine neuron subpopulations. The two STRING networks are based on 23 DE genes highly expressed in SNc (A) and 51 DE genes highly expressed in VTA (B). Different color edges represent the interactions by curated databases, experiment or prediction. The nodes of the two networks are grouped using dash line by using MCL clustering, in which the nodes with solid edges are from subnetworks. (C) The activity of the DEGs present in the SNc and VTA networks highlight both possible beneficial functions as well as functions that could render neurons susceptible, which were predominant in the resilient versus vulnerable dopamine neuron subpopulations. Proposed enriched functions include e.g. neuronal survival, regulation of mitochondrial stability, catabolism of dopamine, regulation of resting membrane potential, extracellular matrix modulation and regulation of cytoskeleton and synapse integrity, G protein signaling and calcium uptake.

### The stable DEGs identified in control tissues also define SNc and VTA subpopulations in PD

We conducted LCM-seq on PD patient tissues to understand if the stable DEGs identified in control tissues would still define SNc and VTA dopamine neurons in end-stage disease. Hierarchical clustering of SNc and VTA PD samples using the stable DEGs separated the majority of samples into the expected subtypes, only one sample out of each group misclassified using this approach. This indicates that the stable genes still define the uniqueness of these two dopamine neuron subpopulations in disease (Fig. 4A). Analysis of individual DEGs showed that *SEZ6, ATP2A3, CBLN1* and *RGS16* maintained a preferential expression in SNc versus VTA dopamine neurons also in PD, although the expression was lower in PD than control SNc (Fig 4B). Similarly, *LY6H, MMP17, EN2, PCSK2, FXYD6* and *PEG3*, defined the VTA subclass of dopamine neurons also in PD, but were also in general lower in PD (Fig 4C). Thus, the identified markers can be used to study the two subpopulations both in health and PD. However, our small PD cohort indicates that the markers may be affected by the disease process. Therefore, we analysed the expression of the stable DEGs in a larger cohort of PD samples where SNc dopamine neuron gene expression was analysed in health and PD [50]. This analysis demonstrated that two stable SNc DEGs, *SLIT1* and *ATP2A3*, were present as significantly lower levels in PD (Fig. 4D).

**Figure 4.**
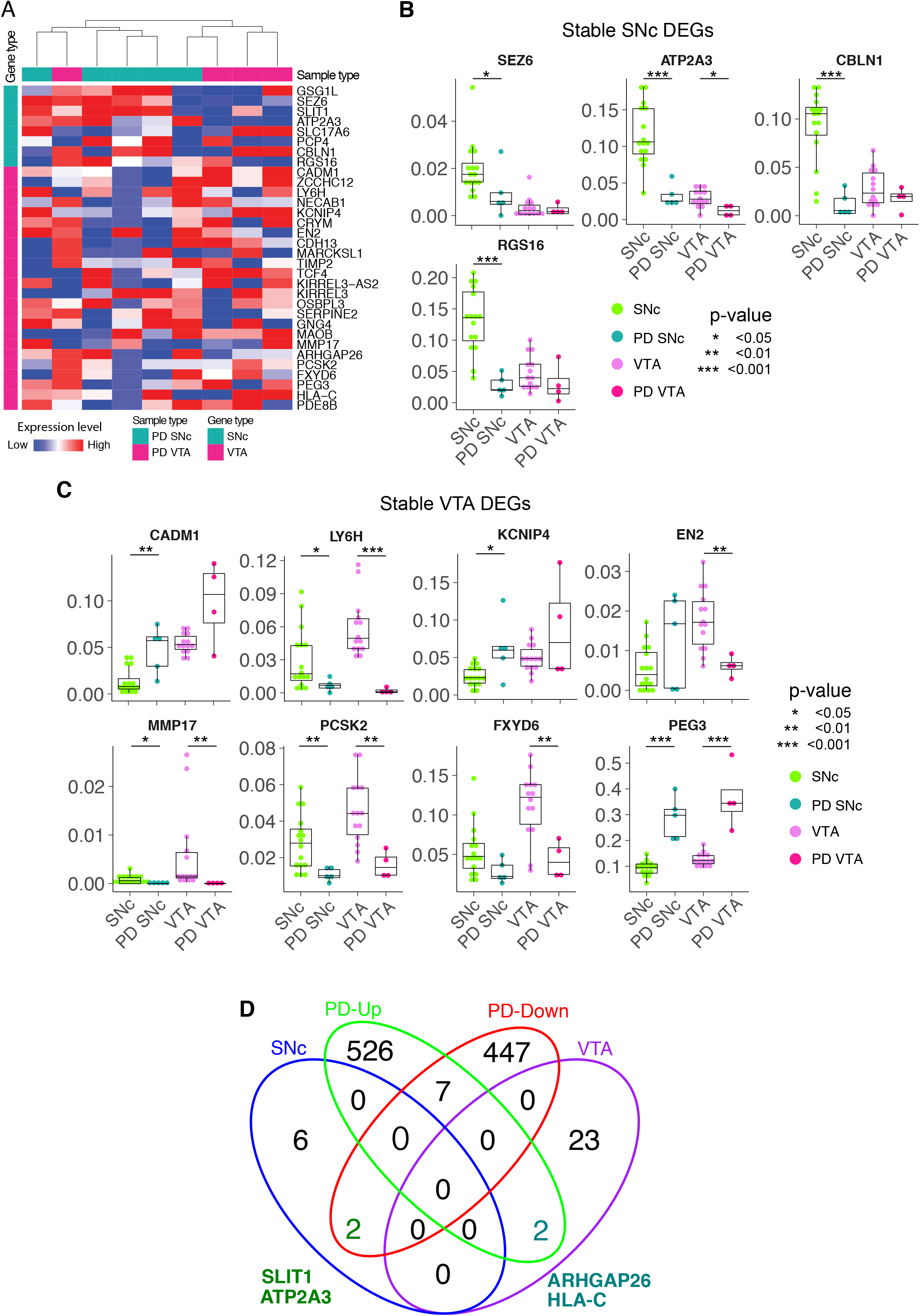
The stable DEGs identified in control tissue separates SNc and VTA from Parkinson’s disease patients. (A) The stable DE genes hierarchically separate SNc and VTA from both normal and Parkinson’s disease (PD) patients. (B, C) The differential expressions of stable DE genes between normal and PD patients in SNc and VTA are shown. (D) Venn diagram finds two common genes, SLIT1 and ATP2A3, between SNc stable and down-regulated genes in PD SNc.

Furthermore, two stable VTA DEGs, ARHGAP26 and HLA-C, were upregulated in the SNc of the large PD cohort (Fig. 4D). The marked decrease of *SLIT1* and *ATP2A3*, and the increase of ARHGAP26 and HLA-C, is a novel PD signature which could be further explored to evaluate neuronal resilience and vulnerability.

## DISCUSSION

The selective vulnerability of SNc dopamine neurons to PD, and the relative resilience of VTA dopamine neurons, has encouraged the field to investigate the molecular signature of these two neuron subpopulations. When we analysed existing data sets [19,11,18,6,45,30,39], we identified large discrepancies in the reported SNc or VTA enriched genes across different studies. This could result from multiple factors, including small sample sizes and variability between subjects, which is recognized to be a major confounding factor in human studies [35]. This prompted us to conduct a large focused study on adult human midbrain dopamine neurons using LCM-seq [38]. We consequently constructed a comprehensive LCM-seq dataset, isolating single SNc or VTA dopamine neurons from *post-mortem* tissues of total 18 individuals, the largest collection of human dopamine neurons, aiming to reveal robust molecular signatures to distinguish the two subpopulations.

Using an iterative bootstrapping without replacement coupled with DESeq2 (available at https://github.com/shanglicheng/BootstrappingWithoutReplacement), and a strict selection criteria (here, a 30% threshold for stable classification) we identify 33 of the most stable DEGs. Among these, 25 of the genes define VTA identity, while eight define SNc identity, which together accurately classify LCM-seq samples from our previous (three females), and current (18 males) subject cohorts. We confirm the utility of our bootstrapping approach on a larger published mouse single cell data set and show that identified DEGs there could correctly classify SNc and VTA dopamine neurons.

Using our approach we also identify a minimal sample size required to identify human stable genes, which for our cohort was an N=8. The sample size may of course vary depending on the specifics of the cohort and the similarity of the subpopulations to be compared. However, our approach clearly demonstrates that to identify lineage-specific markers between any two highly related cellular subpopulations it is of utter importance to determine the sample size and to use a sufficiently large cohort size. Such considerations also apply to studies comparing, for example, healthy and diseased dopamine neurons that may exhibit potentially subtle pathological changes.

The identified stable DEGs highlighted that VTA and SNc dopamine neurons display differences in several important functions such as cytoskeletal regulation, extracellular matrix modulation, synapse integrity, mitochondrial stability, regulation of apoptosis and neuronal survival. The VTA-predominant transcript *SERPINE2* (Glia-derived nexin) is a serine protease inhibitor which can promote neurite extension by inhibiting thrombin, and which appears downregulated in Alzheimer’s disease [10]. Serpine2 promotes biogenesis of secretory granule which is required for neuropeptide sorting, processing and secretion [27]. *CDH13*, another VTA-specific transcript, encodes for adhesion protein 13, which together with other family members as *CDH9* and *CDH15* are linked to neuropsychiatric disorders [47]. Cdh13 can regulate neuronal migration and also has an effect on axonal outgrowth as demonstrated in the serotonergic system [16]. The VTA-predominant gene Engrailed-2 (EN2) is a transcription factor known to promote survival of dopamine neurons by inducing survival gene expression and by protecting neurons from oxidative stress and blocking mitochondrial instability [1,2,48]. The higher level of EN-2 in VTA compared to SNc neurons could in part explain the relative resilience of these neurons to PD. The stable DEGs we identified here may be highly relevant to induce resistance or model disease as previously attempted in rodents [11,45]. Several of the human stable genes (or related family members) e.g. *GSG1L, ATP2A3, SLC17A6, SLIT1, RGS16, KCNIP1, CDH13, TCF12, OSBPL1A, OSBPL10, GNG7, ARHGAP18, ARHGAP24, PCSK5, PEG3, HLA-DOA, HLA-DRA, HLA-DRB1* and *PDE8B* are dysregulated in PD [8,7,51] and/or are represented in PD datasets from genome wide association studies (GWASdb SNP-Disease Associations dataset, http://amp.pharm.mssm.edu. Interestingly, mice lacking *Rgs6*, a related family member of the human SNc stable gene *RGS16*, develop specific degeneration and cell loss of SNc dopamine neurons at the age of 12 months [6]. Loss of the SNc stable gene *Cplx1* results in a compromised nigrostriatal pathway in knockout mice [23]. Moreover, mutations in the human SNc stable gene *SEZ6* have been implicated in diseases such as Alzheimer’s [26,42], childhood-onset schizophrenia [3], epilepsy and febrile seizures [60,37]. CALBINDIN 1 (CALB1) is often used as a marker unique to VTA dopamine neurons. The rank of *CALB1* on the VTA list was just below the 30% frequency threshold for the “stable gene” classification (Fig. 2E). However, while CALB1 is present in the majority of VTA dopamine neurons it is also present in a selection of SNc dopamine neurons [43] and thus it is not surprising that it did not make it onto the stable gene list. Notably, it may be a general marker of resilient dopamine neurons as CALB1^+^ neurons in the SNc show relative sparing in Parkinson’s disease [59].

Analysis of stable DEG expression in PD material showed that SEZ6, ATP2A3, CBLN1 and RGS16 maintained preferential expression in SNc versus VTA dopamine neurons also in disease. Similarly, LY6H, MMP17, EN2, PCSK2, FXYD6 and PEG3, defined the VTA subclass of dopamine neurons also in PD. Analysis of the stable DEGs identified in control brains in a larger cohort of PD samples where SNc dopamine neuron gene expression was analysed demonstrated that two genes, SLIT1 and ATP2A3, out of the eight stable SNc DEGs were dysregulated in PD. This could indicate that these two markers are mainly expressed in the most vulnerable SNc dopamine neurons that are no longer present in end-stage PD patient tissues. However, it is also possible that these two genes are down-regulated in general in all SNc neurons. Future single cell analysis of human dopamine neurons throughout disease progression in PD could aid in discriminating between these two possible scenarios. Nonetheless the marked downregulation of these two markers in PD can be used to distinguish disease-afflicted from healthy SNc dopamine neurons. SLIT1 appears to block neurite extension of dopamine neurons [32]. The loss of SLIT1 may be a compensatory response of remaining cells to allow for neurite growth during disease. Notably, mutant PD-causative forms of LRRK2 induce dystrophic neurites and can also decrease the number of neurites [31,34], indicating that it would be beneficial for dopamine neurons to counteract such processes by modulating the transcriptome accordingly to promote neurite extension. Alternatively, it is possible that the SNc neurons that had high levels of SLIT1 were lost earlier in the PD process due to their inability to modulate neurite extension. This would parallel the situation in amyotrophic lateral sclerosis where motor neurons having high levels of the growth repellant factor EPHA4 are the neurons that are unable to sprout and reconnect with muscle targets and which are consequently lost first in disease [56]. It would be feasible to distinguish between these two possible scenarios using single cell RNA sequencing from *post-mortem* PD tissues from different disease stages.

The lower levels of *ATP2A3*, an ATPase which transports Ca^2+^ across membranes to the endoplasmatic reticulum to maintain a low cytoplasmic Ca^2+^ level, in PD SNc neurons, indicates a deficit in organelle function and Ca^2+^ sequestration. Increased levels of cytoplasmic Ca^2+^ due to lowered ATP2A3 levels could be detrimental to cells and cause degeneration [5]. This data would indicate that remaining SNc neurons have dysfunctions in important cellular processes that need to be tightly regulated by Ca^2+^ levels. The increased level of the stable VTA DEGs, *HLA-C* and *ARHGAP26*, in SNc PD dopamine neurons is very compelling. *ARHGAP26* was recently identified as a potential early, diagnostic biomarker for PD, as it was found upregulated in the blood of PD patients [24]. ARHGAP26 is a Rho GTPase activating protein which is involved in regulating actin-cytoskeleton organization in response to interaction with the extracellular matrix, by mediating RhoA and Cdc42 activity [54]. It would be interesting to study potential structural modifications, resulting from cytoskeletal remodeling, of SNc dopamine neurons in response to PD, and possible effects on their connectome to evaluate if modulating such fundamental processes is part of a protective or detrimental response. HLA-C is a leukocyte antigen that is part of the major histocompatibility complex (MHC)-I, which presents short peptides to the immune system. It has been shown that MHC-I is induced in neurons by factors released from activated microglia, which is a prominent feature of the neuroinflammatory response seen in PD patient tissues. This neuronal MHC-I expression can trigger an antigenic response and cause dopamine neuron death through T-cell mediated cytotoxicity [9]. Thus, an upregulation of *HLA-A* as we see in PD SNc dopamine neurons is likely to be detrimental to the cells.

Regarding cell replacement therapies targeting PD [29,17,01,25,28], there is still an urgent need to optimize the pluripotent stem cell preparations to specifically generate SNc rather than VTA neurons [4,52]. Evaluation of the correct patterning and differentiation of pluripotent cells to midbrain dopamine neurons relies upon gene expression analysis using quantitative real time PCR (qPCR) or global transcriptome approaches such as RNA sequencing [17,4,40,53]. Hence, accurate reference gene signatures of adult human SNc neurons are critical towards further advancements in the regenerative PD field. Our LCM-seq and computational stable gene analysis can therefore serve as a reference describing the transcriptional profile of adult, human SNc and VTA neurons. This will greatly facilitate dopamine neuron replacement efforts, in addition to disease modeling studies using dopamine neurons derived from patient-specific pluripotent cells [36,57].

In summary, using LCM-seq to isolate individual dopamine neurons from SNc and VTA followed by a bootstrapping approach coupled with DESeq2 analysis, we have identified reliable SNc and VTA dopamine neuron markers in human and show that these are relevant also in PD patient tissues. We reveal the smallest human cohort size required to detect such stable DEGs, informing future study designs targeting highly related cellular populations and highlighting that DEGs detected below this cohort size are unreliable. We also demonstrate that a few SNc markers are modulated in PD and could be highly relevant as biomarkers of disease and to understand disease mechanisms further. This human transcriptomic data set, derived from individually isolated dopamine neurons, will thus help further our understanding and modeling of selective neuronal vulnerability and resilience, and serve as a reference for derivation of authentic SNc or VTA dopamine neurons from stem cells.

## ACKNOWLEDGEMENTS

We thank Professor Abdel El Manira and Professor Thomas Perlmann for critical reading of and insightful comments on the manuscript. We thank all Hedlund laboratory members for fruitful discussions. Human *post mortem* tissues were kindly received from the Netherlands Brain Bank (NBB). This work was funded by grants to E.H. from Parkinsonfonden (795/15; 910/16; 991/17; 1095/18; 1192/19); EU Joint Programme for Neurodegenerative Disease (JPND) (529-2014-7500); Swedish Medical Research Council (Vetenskapsrådet) (2016-02112); and NEURO Sweden; by grants to Q.D. from Swedish Research Council (Vetenskapsrådet) (2014-2870); Svenska Sällskapet för Medicinsk Forskning and Jeanssons Stiftelser; Åke Wiberg, Karolinska Institutets Forskningsstiftelser and Stiftelsen; by grants to J.A. from Karolinska Institutets Forskningsstiftelser, Stiftelsen för ålderssjukdomar (2014-2018) and Åhlén-stiftelsen (mA5 h18; Ärende Nr. 193031 h19). J.A. was supported by a postdoctoral fellowship from the Swedish Society for Medical Research and N.K. by a postdoctoral fellowship from Hjärnfonden, Sweden.

## Author contributions

Conceptualization, E.H., Q.D. and J.A.; Methodology and Investigation, J.A., M.C., S.C., N.K., Q.D. and E.H.; Software, Formal Analysis and Visualization, S.C., J.A., and N.K.; Writing-Original Draft, J.A., S.C., Q.D. and E.H.; Writing-review and Editing, J.A., S.C., N.K., M.C., Q.D. and E.H.; Supervision and Project Administration, E.H. and Q.D.; Funding Acquisition, E.H., Q.D. and J.A.

