## Supplemental Figures 1-6, Supplemental Tables 1-3 for "Spatial RNA sequencing identifies robust markers of vulnerable and resistant human midbrain dopamine neurons and their expression in Parkinson’s Disease"

**Table of contents**

Supplemental Figure 1

Supplemental Figure 2

Supplemental Figure 3

Supplemental Figure 4

Supplemental Figure 5

Supplemental Figure 6

Supplemental Table 1

Supplemental Table 2

Supplemental Table 3

*Supplemental Tables 4 to 8. List of DEGs, Sequencing Stats and Stable Genes (combined in 1 excel file with 6 sheets)*

Supplemental Table 4. List of differentially expressed genes from previous transcriptomic studies. Related to Supplemental Figure S1.

Supplemental Table 5. List of experimental variables and sequencing stats for the 43 samples associated with the current dataset (18 male subjects). Related to Figures 1, 2 and Supplemental Figure S2.

Supplemental Table 6. List of differentially expressed genes from the current dataset (18 male subjects, 74 DEGs). Related to Figures 1 and Supplemental Figure S2.

Supplemental Table 7. List of differentially expressed genes from the Nichterwitz's dataset [39] re-analyzed here (3 female subjects, 100 DEGs). Related to Figures 1 and Supplemental Figure S2.

Supplemental Table 8. Full list of human stable genes/DEGs ranked by frequency identified with the bootstrapping approach coupled with DESeq2. See also METHODS section. Related to Figures 1, 2 and Supplemental Figure S3.

#### Supplemental Figure 1

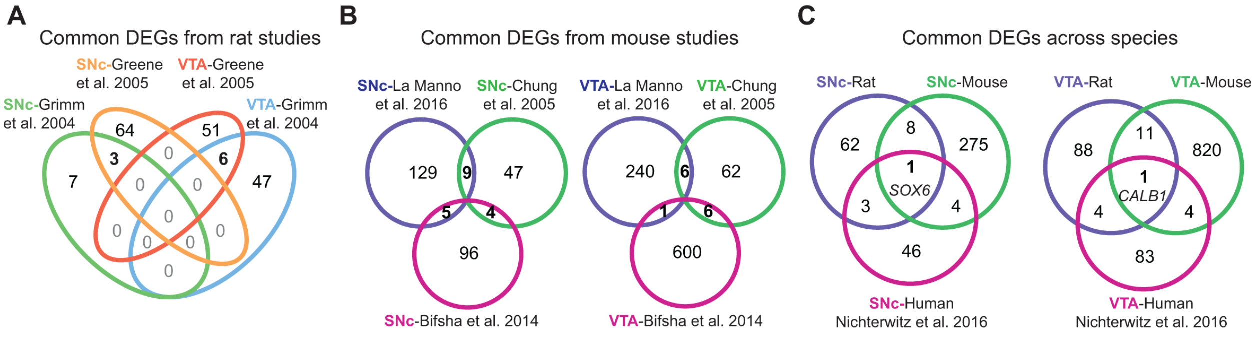


**Supplemental Figure 1.** Common differentially expressed genes across species. Related to Supplemental Tables S1 and S4. (A) Venn diagram showing overlapping DEGs for SNc and VTA identified in previous rat studies [19, 20]. (B) Common mouse DEGs are highlighted for SNc and VTA from three different reports [12,6,30]. Samples from the La Manno dataset [30] were here analyzed with DESeq2, identifying 390 DEGs. (C) Overlapping DEGs between rat, mouse and human [39]. For rat and mouse, all previously reported DEGs were considered despite of the variability within studies as shown in (A) and (B).

### Supplemental Figure 2

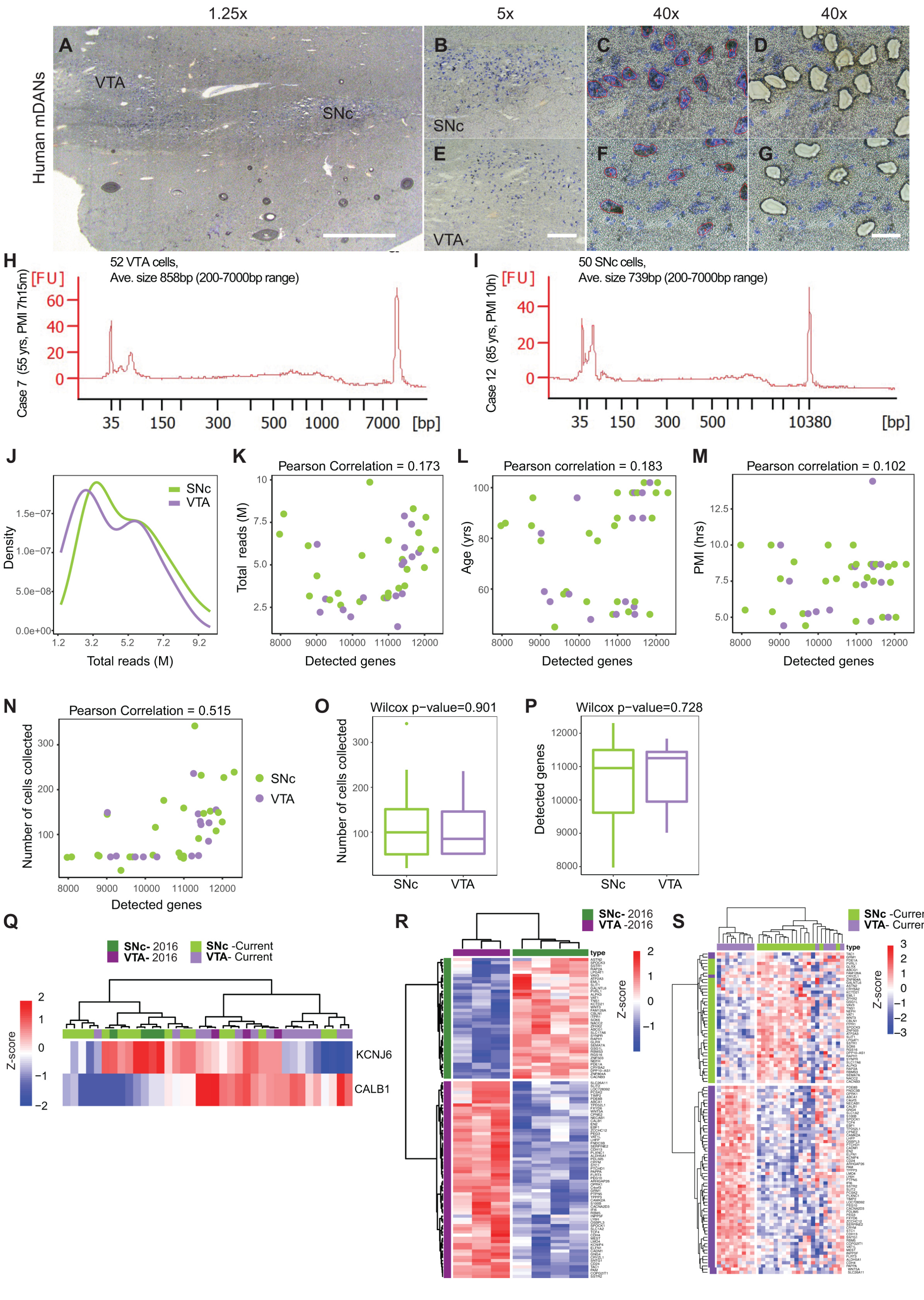

**Supplemental Figure 2.** Quality control of human adult midbrain dopamine neuron samples profiled using LCM-seq. Related to Figure 1 and Supplemental Tables S2, S5, S6 and S7. (A to G) Nissl staining of a representative midbrain section for the isolation of SNc (B,C,D) or VTA (E,F,G) neurons before and after laser capture. (H, I) Representative bioanalyzer profiles of cDNA libraries from VTA (H) and SNc (I) samples. (J-P) The average length size for the cDNA, 700-900 base pairs, was comparable across VTA and SNc. The quality and similarity of samples from both populations was also interrogated based on sequencing stats, including total reads (J) and detected genes (K). This later outcome was also correlated to biological and experimental variables such as the "Age" (L), "Post-mortem interval" (M) and "Cells" collected during the laser capture sessions (N). (O, P) While there was a modest correlation between the number of cells collected and the number of detected genes (N, P=0.515), these variables were not significantly different between SNc and VTA groups. See also Supplemental Table S5. (Q) Hierarchical clustering of VTA and SNc samples using only *KCNJ6* (*GIRK2*) and *CALB1*. (R, S) Samples from the Nichterwitz's study [39] were re-analyzed with DESeq2 (version 1.16.1) which identified 100-DEGs used to represent those samples (P, N=3 subjects) or our current dataset (Q, N=18 subjects). Scale bars: 500 μm in (A), 200 μm in (B, E), 50 μm in (C,D,F,G).

#### Supplemental Figure 3

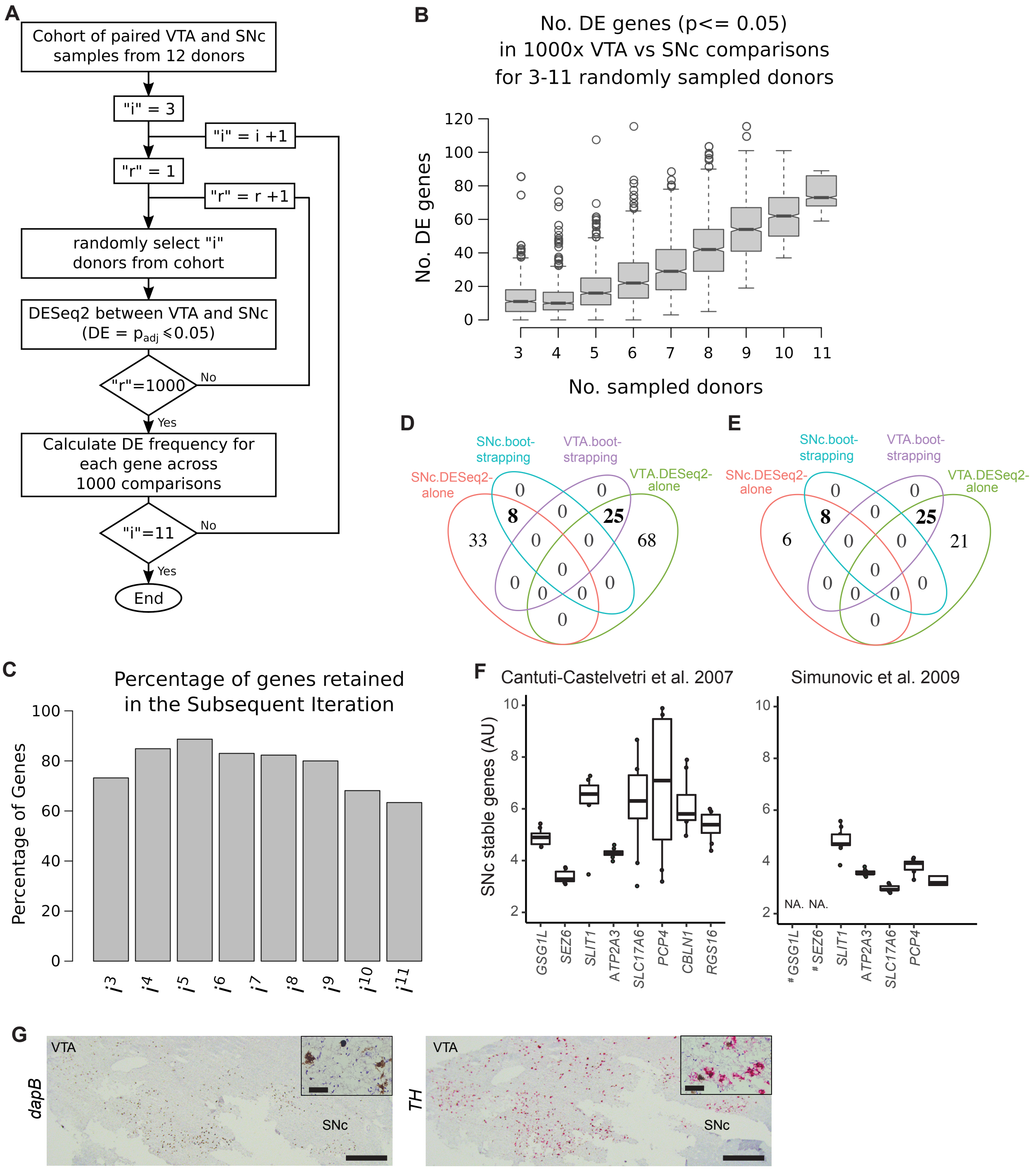


**Supplemental Figure 3.** Outcome of the bootstrapping approach coupled with DESeq2 applied to human samples and validation of SNc stables genes in the context of previous studies. Related to Figures 1, 2 and Supplemental Tables S2 and S8. (A) Flowchart illustrating the computational pipeline for iterative bootstrapping approach, i refers to the sample size of each bootstrapping iteration out of total 12 samples

(3<=i<=11); r refers to iterative times of bootstrapping round in the iteration (r maximum = 1000). (B) Boxplot displaying the total number of unique DEGs between VTA and SNc detected in individual iterations. e.g. over 1000x comparisons in i donors. (C) Percentage of DEGs in a given iteration, that were retained in the subsequent iterations (3-11 subjects). (D, E) To confirm the authenticity of the genes considered stable (33 markers identified through bootstrapping coupled with DESeq2), Venn diagrams were used to assess the overlap with DEGs identified by DESeq2 alone applied to the same 12 subjects (D, adj. *P*-value <0.05; E, adj. *P*-value <0.01). (F) SNc stable genes are expressed in control human SNc dopamine neurons isolated by LCM from previous microarray studies. [#]In the study reported by Simunovic et al. [50], the genes *GSG1L* and *SEZ6* were not included in the array. (G) RNAscope staining of human melanized midbrain sections using a negative control probe against a bacterial gene (*dapB*) and a positive control probe against tyrosine hydroxylase (*TH*). Scale bars: 1mm in (G) with 50 μm in insets.

### Supplemental Figure 4

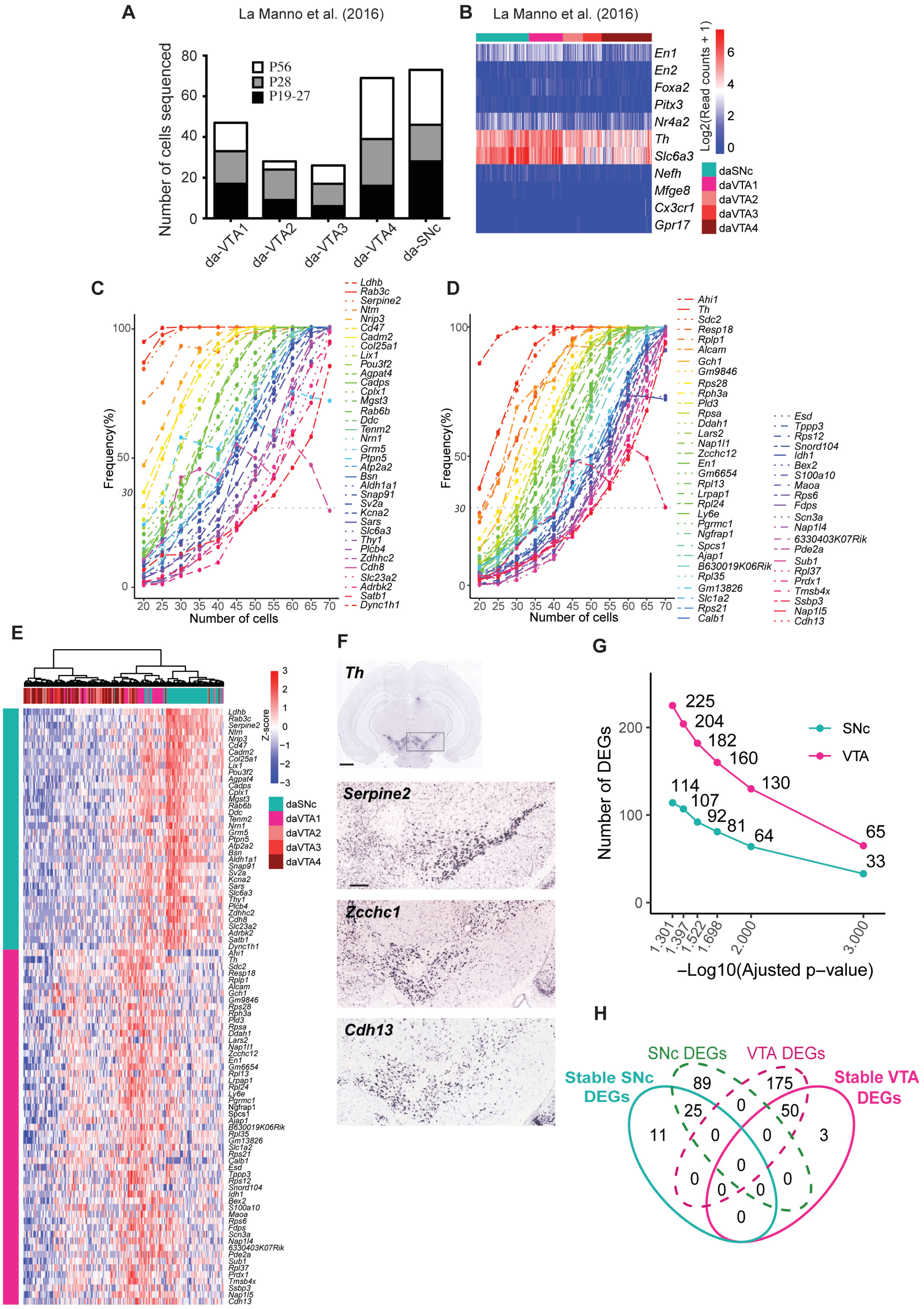

**Supplemental Figure 4.** Validation and outcome of the bootstrapping approach coupled with DESeq2 applied to a published single cell mouse data set. (A) Mouse dataset [30] used to confirm the bootstrapping approach and to identify stable genes in juvenile/adult (postnatal day 19 to 56) midbrain dopamine neurons. The graph shows the number of single cells sequenced for each dopaminergic population. (B) Heat map to confirm the expression of known midbrain dopamine neuron markers and the absence of markers for other neural cells as astrocytes, microglia or oligodendrocytes. (C, D) Identification of SNc (A) and VTA (B) stable genes in the mouse dataset from [30] using the bootstrapping approach coupled with DESeq2 applied to 146 single cells (73 available SNc cells and 73 randomly selected VTA cells). (E) Identified mouse stable genes separate SNc and VTA samples as visualized in the unsupervised clustering. (F) *In situ* hybridization images of coronal midbrain sections of the adult mouse (P56, Allen Brain Atlas) showing RNA expression pattern for *Th*, *Serpine2*, *Zcch12* and *Cdh13*. (G) The number of DEGs in SNc and VTA from [30] identified by DESeq2 as a function of different adjusted P-value cutoffs. (H) Venn diagram showing overlaps among stable SNc DEGs, stable VTA DEGs and DEGs in (G) with adjust an p-value=0.05.

#### Supplemental Figure 5

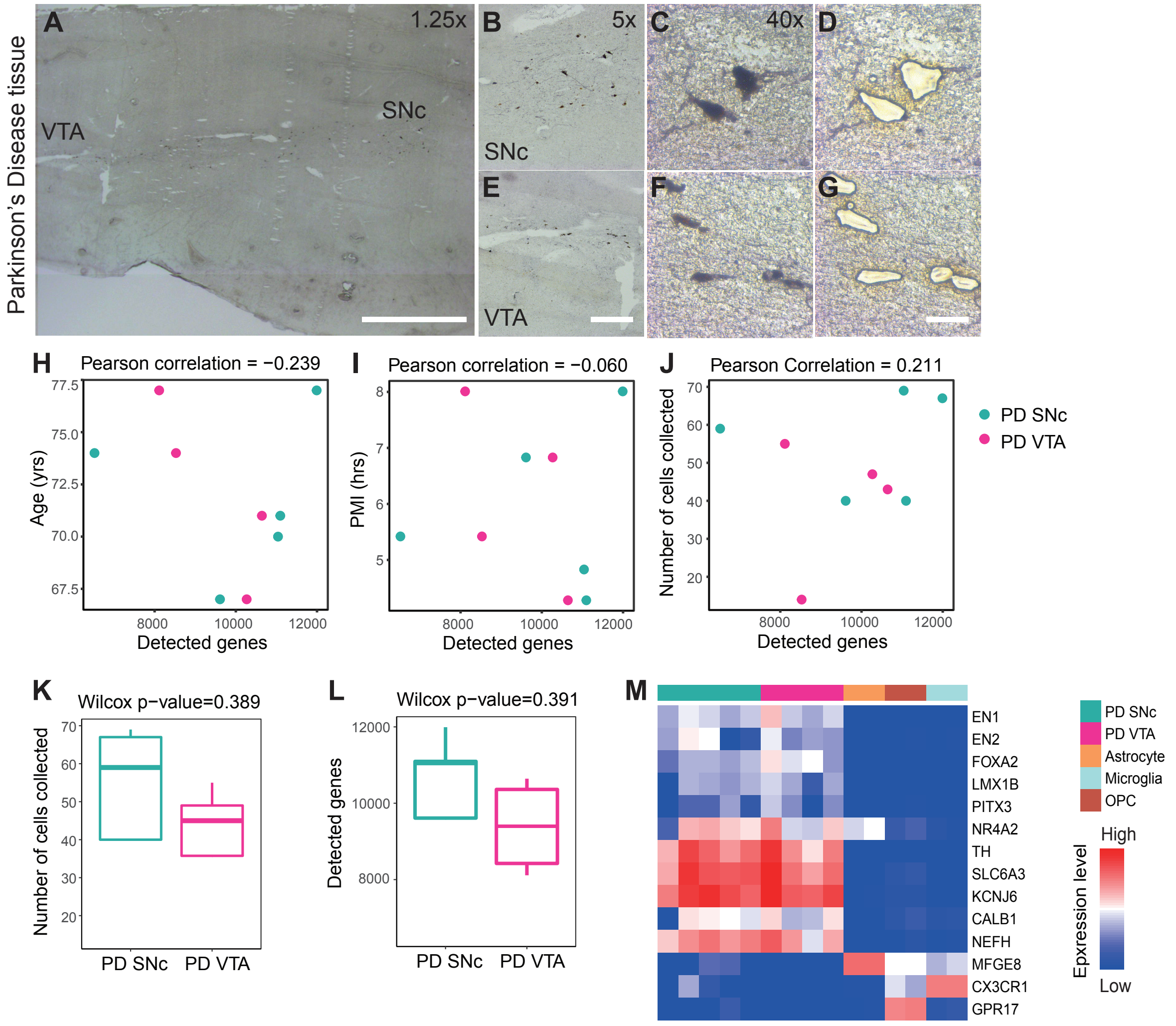


**Supplemental Figure 5.** Quality control of Parkinson's Disease (PD) patient midbrain dopamine neuron samples profiled using LCM-seq. Related to Figure 5 and Supplemental Table S2. (A to G) Nissl staining of a representative midbrain section for the isolation of SNc (B-D) or VTA (E-G) dopamine neurons from PD patient tissues before and after laser capture. The quality and similarity of samples from SNc and VTA populations was interrogated based on sequencing data and the number of detected genes as a function of "Age" (yrs) of the patients (H), "Post-mortem interval, PMI" (hrs) (I) and "The number of cells collected" (J) during the laser capture sessions. None of these variables were significantly different between the SNc and VTA groups. Furthermore, the was no significant difference in the number of cells collected for SNc versus VTA (K) or the number of detected genes for SNc versus VTA PD samples (L). The high sample quality of the PD samples was confirmed by strong expression of the midbrain dopamine neuron markers *EN1/2*, *FOXA2*, *LMX1B*, *PITX3*, *NR4A2*, *TH* and *SLC6A3* (DAT), the pan-neuronal marker neurofilament (*NEFH*), and the lack of astrocyte, microglia or oligodendrocyte precursor marker contamination, as compared to a published study [61].

#### Supplemental Figure 6

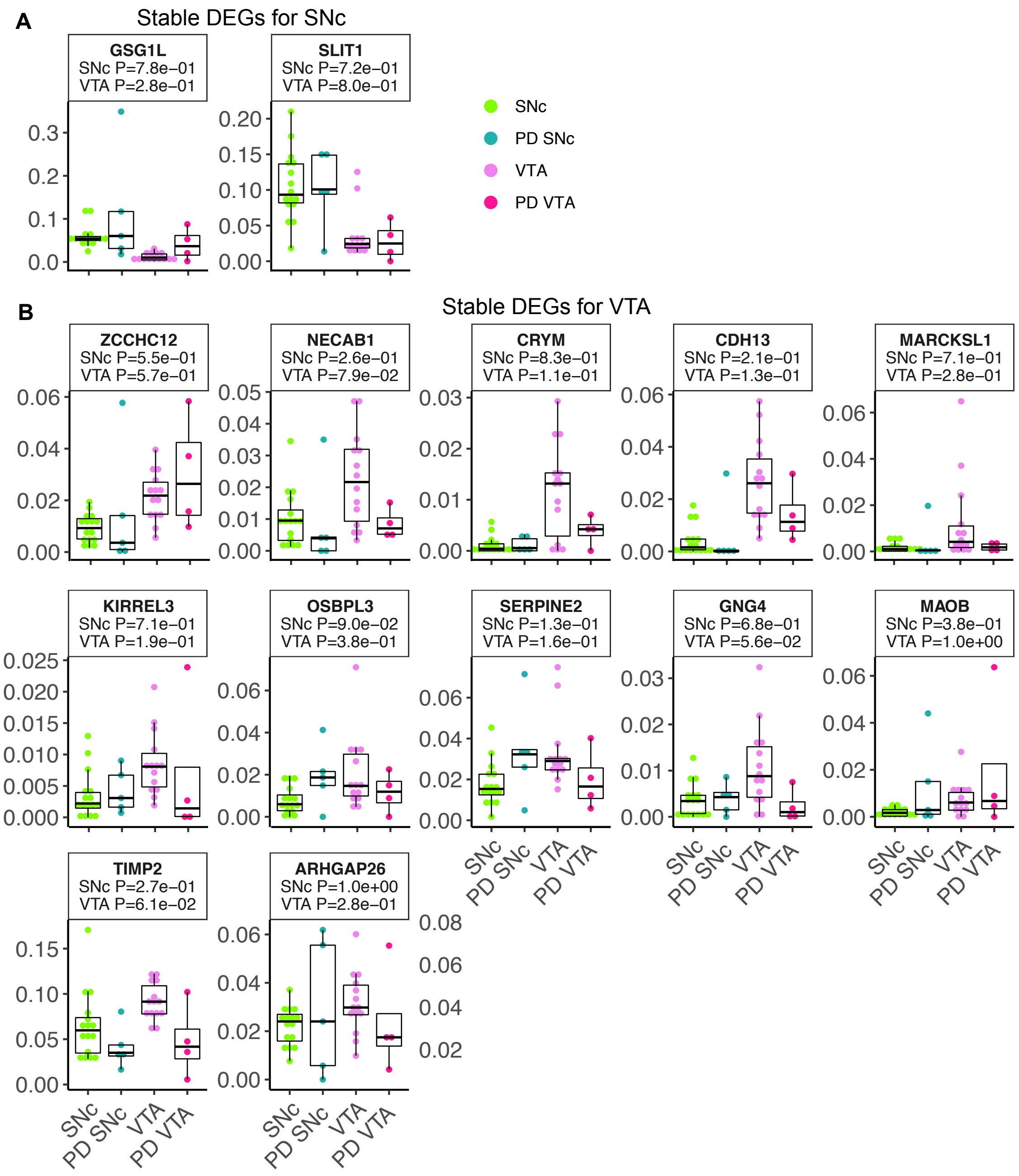


**Supplemental Figure 6.** Boxplots displaying the expression level of the stable DEGs defining SNc and VTA dopamine neurons from both healthy and PD tissues. (A) Two stable DEGs for SNc. (B) Twelve stable DEGs for VTA.

**Supplemental Table 1.** Summary of studies addressing the identity of midbrain dopamine neurons.

| Pubmed ID | Sample origin | Sample size | Method | No of DEGs |
|---|---|---|---|---|
| 15353588 | Sprague–Dawley rats, adult, female, | >4 | qTH stain, LCM bulk, microarray | 66 |
| 15649693 | Lewis rats, adult male | n=8 | qTH stain, LCM bulk, microarray | 124 |
| 15888489 | C57/Bl6 mice, adult | n=5-6 | qTH stain, LCM bulk, microarray | 134 |
| 25437550 | C57/Bl6 mice, P4P4 | mix of three animals | Slc6a3::Cre, FACS-qPCR, single cell array | NA. |
| 25501001 | C57/Bl6 mice, P1-P4 | duplicates (>2) | TH-EGFP, FACS bulk, microarray | 712 |
| 27716510 | CD-1 mice, juvenile – adult | 1-3 brains/time point | FACS and C1 chip, Fluidigm, single cell, RNAseq CD-1, Dat1-Cre/tdTomato) | 390 (calculated here) |
| 17412603 | Human, adult (male and female) | 8 controls | Methylene blue/melanized SNc, LCM bulk, microarray | NA. |
| 19052140 | Human, adult, male and female | 9 controls | melanized SNc, LCM bulk, microarray | NA. |
| 27387371 | Human, adult, female | 3 controls | Histogen, LCM-seq bulk, RNAseq | 111 |
| Current study | Human, adult (male and female) | 18 controls<br>5 PD patients | Histogen, LCM-seq bulk, RNAseq | 74 |

NA. Not applicable.
qTH. Quick tyrosine hydroxylase antibody staining.
FACS. Fluorescence-activated cell sorting
qPCR. Quantitative polymerase chain reaction
LCM. Laser capture microdissection.
RNAseq. RNA sequencing

**Supplemental Table 2.** Clinical information of subjects used for LCM-seq.

| | **Gender** | **Age** | **Post mortem time** | **Cause of death** | **Brain Bank** | **Targeted cells** |
|---|---|---|---|---|---|---|
| 1 | M | 45 | 8h 50m | Intestinal metastasis | NBB | SNc |
| 2 | M | 48 | 5h 30m | Euthanasia, diabetes | NBB | VTA |
| 3 | M | 50 | 8h 30m | Cardiac arrest | NBB | SNc and VTA |
| 4 | M | 51 | 7h 45m | Suicide, self-withering | NBB | SNc and VTA |
| 5 | M | 53 | 14h 25m | Heart failure | NBB | VTA |
| 6 | M | 55 | 7h 30m | Euthanasia, cancer | NBB | SNc and VTA |
| 7 | M | 55 | 7h 15m | Thrombosis | NBB | SNc and VTA |
| 8 | M | 58 | 5h 15m | Thrombosis | NBB | SNc and VTA |
| 9 | M | 59 | 4h 25m | Myocardial infarction | NBB | SNc and VTA |
| 10 | M | 79 | 7h 40m | Pneumonia and sepsis | NBB | SNc |
| 11 | M | 82 | 10h | Pleuritis, cachexia | NBB | SNc and VTA |
| 12 | M | 85 | 10h | Prostate cancer, asthma | NBB | SNc |
| 13 | M | 86 | 5h 30m | Cancer, cachexia | NBB | SNc |
| 14 | M | 88 | 4h 43m | Intestinal bleeding | NBB | SNc and VTA |
| 15 | M | 88 | 7h 25m | Euthanasia, cancer | NBB | SNc and VTA |
| 16 | M | 96 | 5h 23m | Pneumonia | NBB | SNc and VTA |
| 17 | M | 98 | 8h 40m | Cardiac tamponade | NBB | SNc and VTA |
| 18 | M | 102 | 5h | Ileus | NBB | SNc and VTA |
| 19 | M | 67 | 6h50m | PD, Pneumonia and dehydration | NBB | SNc and VTA |
| 20 | M | 70 | 4h50m | PD, Cachexia and dehydration | NBB | SNc |
| 21 | M | 71 | 4h17m | PD, Urinary tract infection | NBB | SNc and VTA |
| 22 | M | 74 | 5h25m | PD, Euthanasia | NBB | SNc and VTA |
| 23 | F | 77 | 8h07m | PD, Pneumonia | NBB | SNc and VTA |

M, male
F, female
SNc, substantia nigra compacta
VTA, ventral tegmental area
PD, Parkinson disease (All cases were PD with dementia).
NBB, Netherlands Brain Bank, www.brainbank.nl

**Supplemental Table 3.** Clinical information of subjects used for RNAscope.

| Case | Gender | Age | Post mortem time | Cause of death | Brain Bank |
|---|---|---|---|---|---|
| 1 | M | 48 | 5h 30m | Euthanasia | NBB |
| 2 | M | 50 | 8h 30m | Cardiac arrest | NBB |
| 3 | M | 55 | 7h 15m | Intestinal ischemia | NBB |
| 4 | M | 82 | 10h 00m | Pleuritis carcinomatosis, cachexia | NBB |
| 5 | M | 88 | 4h 43m | Gastrointestinal bleeding | NBB |
| 6 | M | 98 | 8h 40m | Cardiac tamponade, atherosclerotic aorta | NBB |

M, male
NBB, Netherlands Brain Bank

**Supplemental References (all within the reference list of the article)**
